## Supplementary information for "A non-image-forming visual circuit mediates the innate fear of heights"

**Supplementary Table 1 Coordinates and virus used for stereotaxic injection.**

| Brain region | Coordinates (mm) | Volumes (µl) |
|---|---|---|
| LPMR | *AP*: -2.06 *ML:* ± 0.95 *DV*: -2.60 | 0.20 |
| SC | *AP*: -4.00 *ML:* ± 0.45 *DV*: -1.80 | 0.40 |
| V1 | *AP*: -3.50 *ML:* ± 2.30 *DV*: -1.40 | 0.40 |
| vLGN | *AP*: -2.35 *ML:* ± 2.50 *DV*: -3.60 | 0.15 |
| l/vlPAG | *AP*: -4.25 *ML:* ± 0.60 *DV*: -2.95 | 0.20 |
| BLA | *AP*: -1.46 *ML:* ± 3.50 *DV*: -4.75 | 0.20 |
| CeA | *AP*: -1.22 *ML:* ± 2.90 *DV*: -4.75 | 0.15 |

| Virus | Supplier/Lot. number | Titer (µg $ml^{-1}$) |
|---|---|---|
| AAV2/9-*hSyn*-hM4Di-EGFP | BrainVTA, PT-0153 | $5.15\times10^{12}$ |
| AAV2/Retro-*hSyn*-Cre-EGFP | Braincase, BC-0160 | $5.00\times10^{12}$ |
| AAV2/9-*hSyn*-DIO-hM4Di-mCherry | Braincase, BC-0153 | $5.06\times10^{12}$ |
| AAV2/9-*hSyn*-DIO-hM3Dq-mCherry | Braincase, BC-0143 | $5.47\times10^{12}$ |
| AAV2/9-*hSyn*-DIO-mCherry | Braincase, BC-0025 | $5.65\times10^{12}$ |
| AAV2/9-Vglut2-hM4Di-EGFP | BrainVTA, PT-3883 | $5.31\times10^{12}$ |
| AAV2/9-*hSyn*-DIO-GCaMP6s | Braincase, BC-0238 | $3.02\times10^{12}$ |

**Supplementary Table 2 Antibodies used for immunofluorescent staining.**

| Antibody | Company | Catalog number | Dilution ratio |
|---|---|---|---|
| Anti-c-fos (Rabbit) | CST | 2250S-100 μl | 1:750 |
| Anti-c-fos (Mouse) | Thermo Fisher | MA1-21190 | 1:1000 |
| Anti-GABA (Rabbit) | Sigma | A2052 | 1:500 |
| Anti-Parvalbumin (Rabbit) | Abcam | ab11427 | 1:500 |
| IgG H&L (Alexa Fluor® 647) (Goat anti-rabbit) | CST | 4414S | 1:1000 |
| IgG H&L (Alexa Fluor® 488) (Goat anti-rabbit) | Thermo Fisher | A-11029 | 1:1000 |
| IgG H&L (Alexa Fluor® 546) (Goat anti-rabbit) | Thermo Fisher | A-21089 | 1:1000 |

**Supplementary Figure 1**

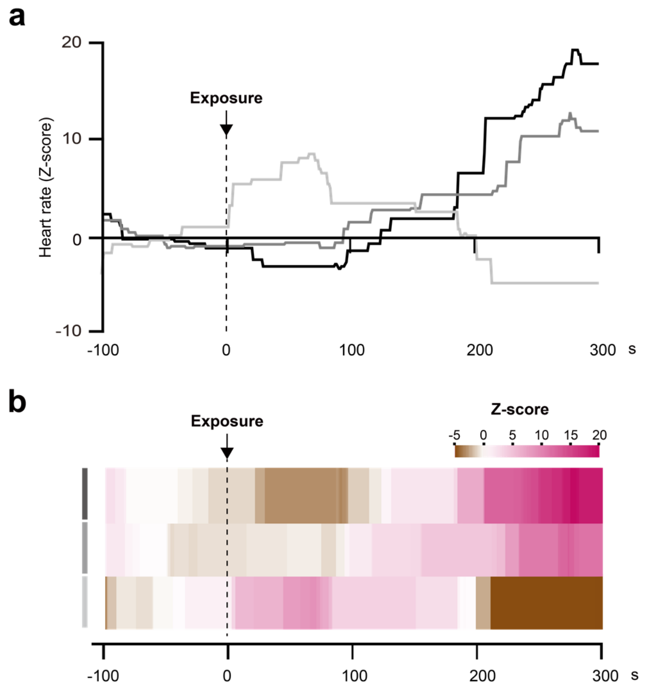

**Fig. S1 The heart rate of mice increases upon height exposure. a** Heart rate (HR) fluctuation over time (s, second) before and after height exposure. Z-score ((observed HR-baseline HR)/standard deviation of baseline) is used to illustrate HR changes. **b** Heatmaps of HR change over time based on the data from (**a**). Results of three mouse individuals are shown.

**Supplementary Figure 2**

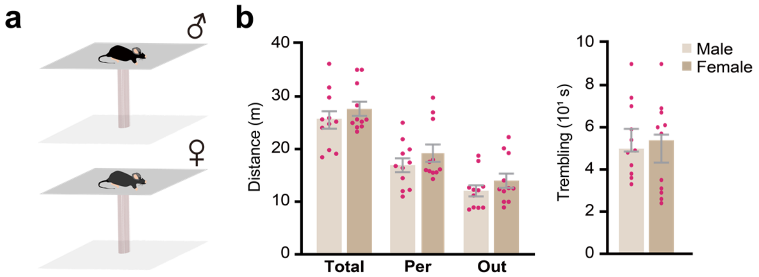

**Fig. S2 Male and female mice have similar anxiety level in the OHP test. a** Schematic diagram of male and female mice in the OHP test. **b** No significant differences are identified between male and female mice in all parameters analyzed. Data are presented as the mean ± S.E.M.

### Supplementary Figure 3

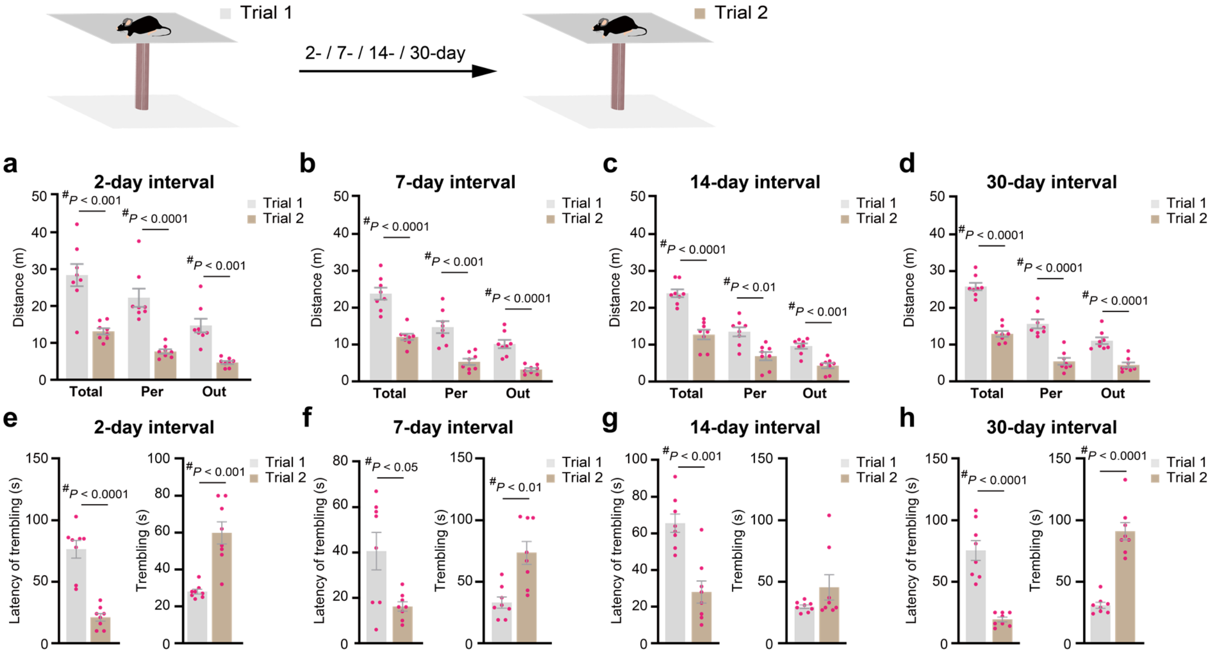

**Fig. S3 Facilitation of fear of heights upon re-exposure of mice to height threat with different time intervals. a-d** Comparison of the locomotion distances of mice at different regions of the OHP in two trials with different intervals. **e-h** Comparison of the latency of trembling initiation and the total duration of trembling on the OHP in two trials with different intervals. Data are presented as the mean ± S.E.M. $^{\#}p$ <0.05, two-tailed Student's *t*-test.

### Supplementary Figure 4

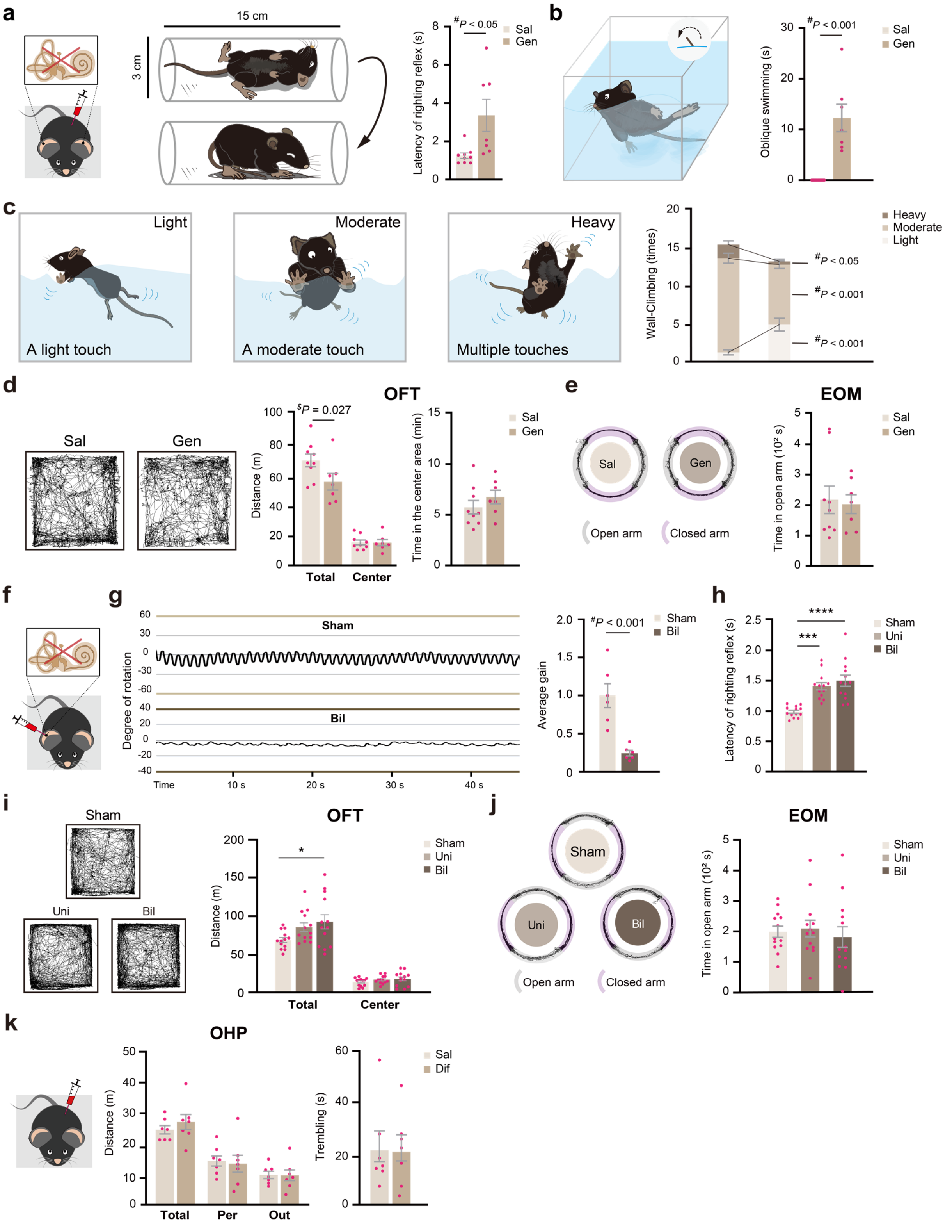

**Fig. S4 Vestibular input is dispensable to fear of heights. a-c** Results of the latency of righting reflex (**a**) and swimming test (**b**, **c**) for control and gentamicin-treated mice. The total time of oblique swimming (**b**) and the total counts for light, moderate, and heavy wall-climbing events (**c**) are shown. **d** Total locomotion distance and total time spent in the center of the open field arena of control and gentamicin-treated mice in the open field test (OFT). **e** The open arm exploration time of control and gentamicin-treated mice in the elevated O-maze test (EOM). **f-j** Results of the vestibulo-ocular reflex (VOR, **f, g**), righting reflex (**h**), OFT (**i**), and EOM (**j**) for mice received the unilateral (Uni) or bilateral (Bil) intratympanic application of sodium arsanilate. Time series plot of the eye movement and average gain values (Eye movement velocity/Head movement velocity) of sham and bilaterally treated mice are shown in **f** and **g**. (**k**) Results of OHP test for difenidol-treated mice. Data are presented as the mean ± S.E.M. with a One-Way ANOVA and a one-tailed ($^{\$}$) or a two-tailed ($^{\#}$) Student's *t*-tests.

**Supplementary Figure 5**

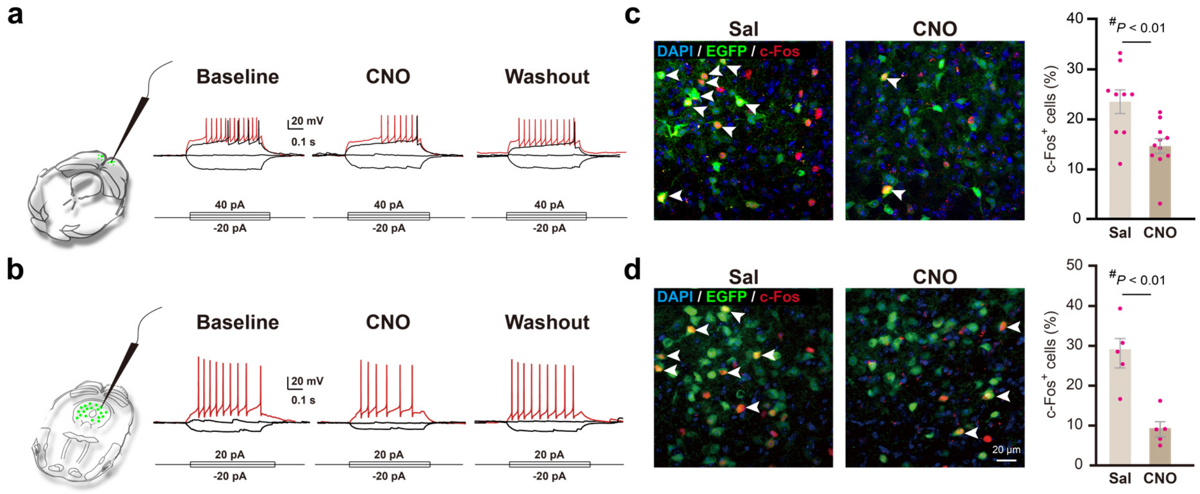

**Fig. S5 Validation of the effectiveness of chemogenetic inhibition. a, b** Validation of the effectiveness of chemogenetic inhibition of neuronal activities by whole-cell patch-clamp recording in brain slices from mice infected with the inhibitory virus vector hM4Di (AAV2/9-*hSyn*-hM4Di-EGFP) in upper layers of SC (**a**) or the l/vlPAG (**b**). Fluorescent cells that express the hM4Di vector were recorded and their firing before, during, and after washing out CNO treatment are shown. **c, d** Immunofluorescence staining of c-Fos. Coronal brain sections are from mice injected the hM4Di vector in upper layers of SC (**c**) or the l/vlPAG (**d**), treated with or without CNO. The percentage of $EGFP^+$ cells that are $Fos^+$ with or without CNO treatment are shown. Data were analyzed using a two-tailed Student's *t*-test ($^{\#}$) and are presented as the mean ± S.E.M.

### Supplementary Figure 6

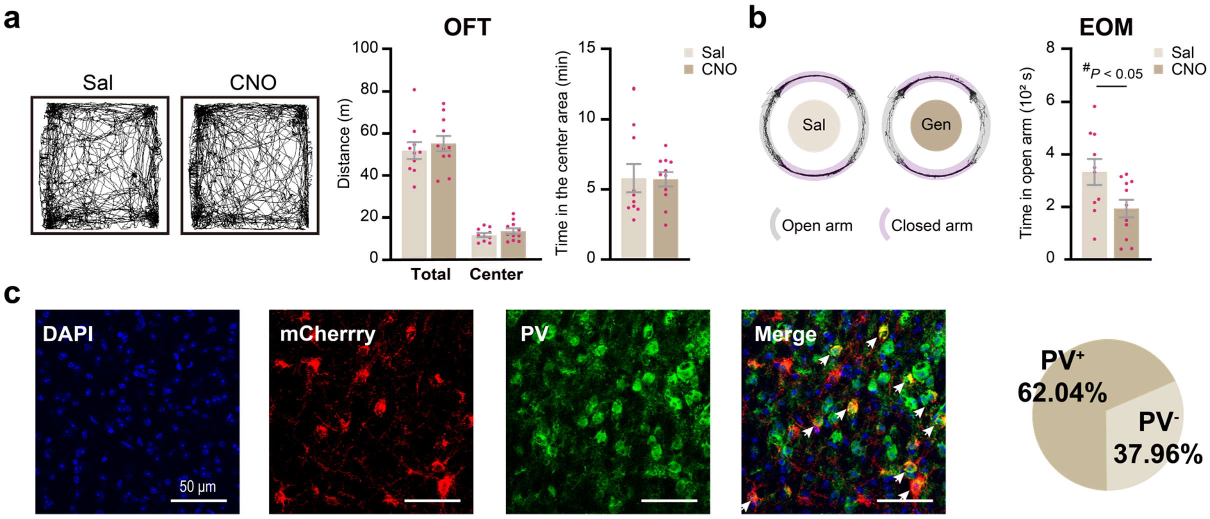

**Fig. S6 Characterization of potential role of the SC. a**, **b** Effects of chemogenetic inhibition of pan-neuronal activities of SC (AAV2/9-*hSyn*-hM4Di-EGFP) on the locomotion and anxiety level of mice in the OFT (**a**) and EOM (**b**). Data are presented as the mean ± S.E.M. $^{\#}p < 0.05$, a two-tailed Student's *t*-test. **c** Parvalbumin-positive ($PV^{+}$) neurons constitute the majority of projections from the SC to LPMR. Retrograde tracing was achieved by injecting AAV2/9-*hSyn*-DIO-hM4Di-mCherry (red) into the SC and AAV2/Retro-*hSyn*-Cre-EGFP into LPMR. SC brain slices were stained for PV (green, pseudo color).

### Supplementary Figure 7

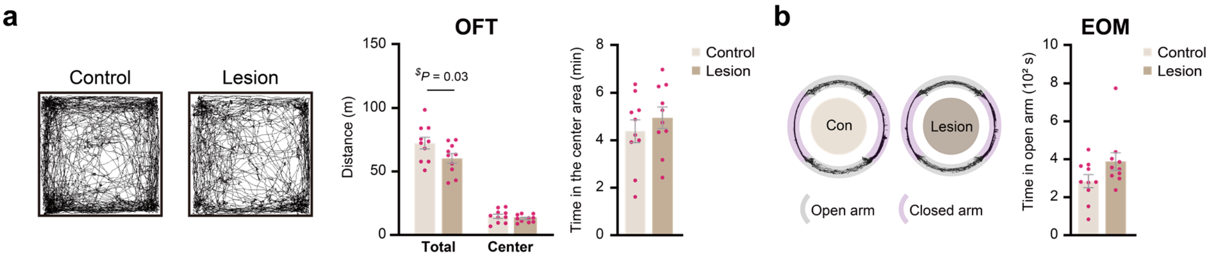

**Fig. S7 V1 lesion does not affect the anxiety level of mice. a, b** Effects of V1 excision on mouse behaviors in the OFT (**a**) and EOM (**b**). Data are presented as mean ± S.E.M. $^{\$}p < 0.05$, a one-tailed Student's *t*-test.

**Supplementary Figure 8**

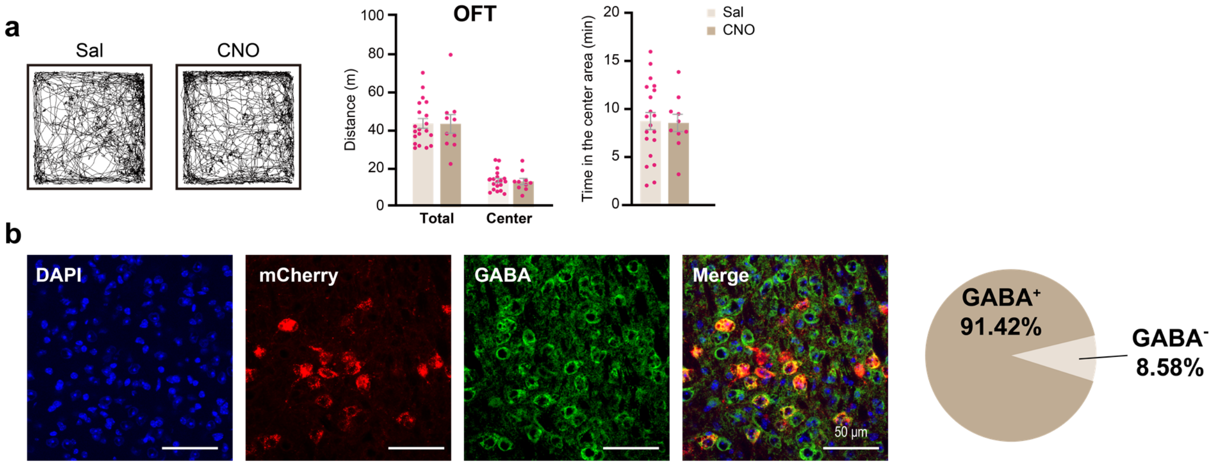

**Fig. S8 Role of vLGN in the regulation of mouse behaviors. a** Effect of pan-neuronal inhibition of vLGN on mouse behaviors in the OFT. **b** Immunofluorescence fate-mapping of vLGN-to-l/vlPAG projecting neurons using retrograde tracing. Retrograde tracing was achieved by injecting AAV2/Retro-*hSyn*-Cre-EGFP into l/vlPAG and AAV2/9-*hSyn*-DIO-hM4Di-mCherry (red) into vLGN. The vLGN brain slices were stained with the antibody against GABA (pseudo colored green). The pie chart shows the the proportion of $GABA^+$ cells relative to the total population of $mCherry^+$ cells. Data are presented as mean ± S.E.M.

**Supplementary Figure 9**

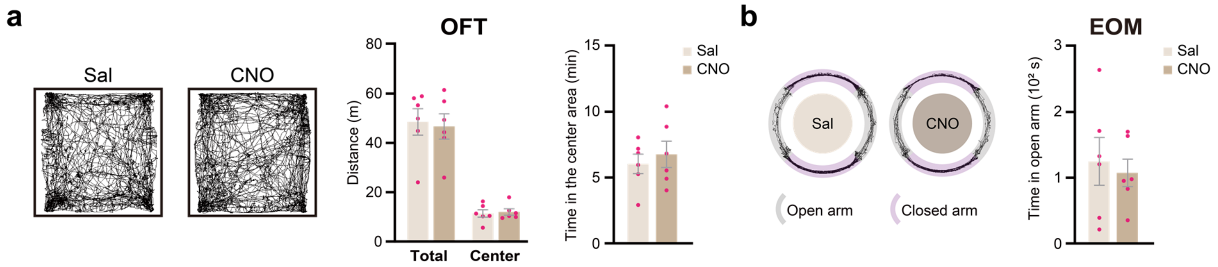

**Fig. S9 Inhibition of l/vlPAG causes no significant changes in the general anxiety level.** **a, b** Effects of chemogenetic inhibition of pan-neuronal activities of l/vlPAG on mouse behavior in the OFT (**a**) and EOM (**b**). Data are presented as the mean ± S.E.M.

### Supplementary Figure 10

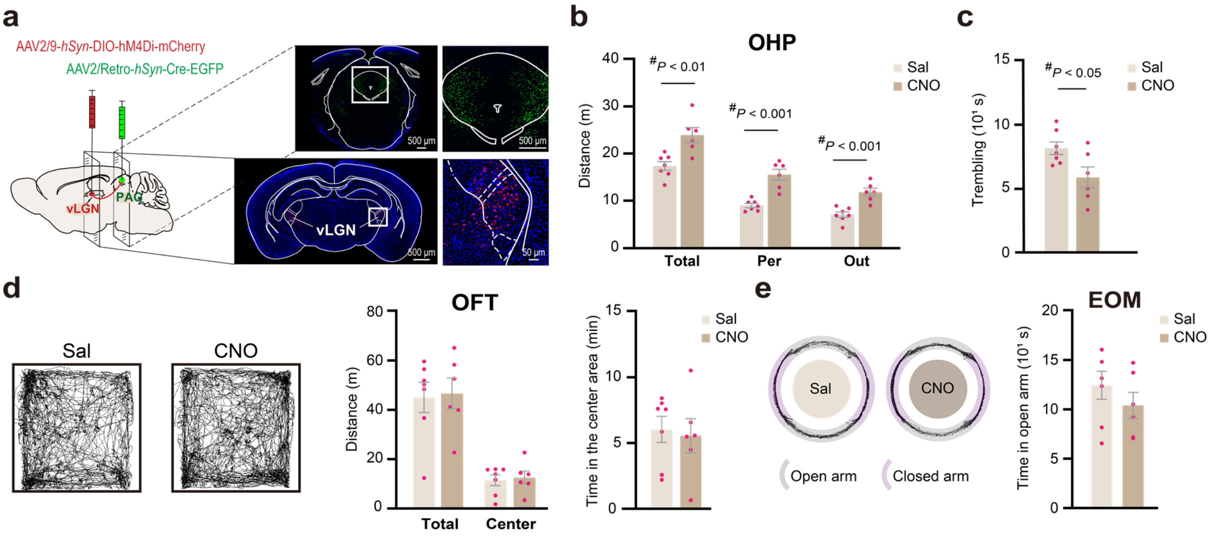

**Fig. S10 Inhibition of the vLGN-PAG projecting neurons reduces fear of height. a-c** Effects of chemogenetic inhibition of neurons that project from the vLGN to the l/vPAG on the fear of heights. **d, e** Effects of chemogenetic inhibition of neurons that project from vLGN to the l/vlPAG on mouse behaviors in the OFT and EOM. Data are presented as the mean ± S.E.M. $^{\#}p < 0.05$, a two-tailed Student's *t*-test.

**Supplementary Figure 11**

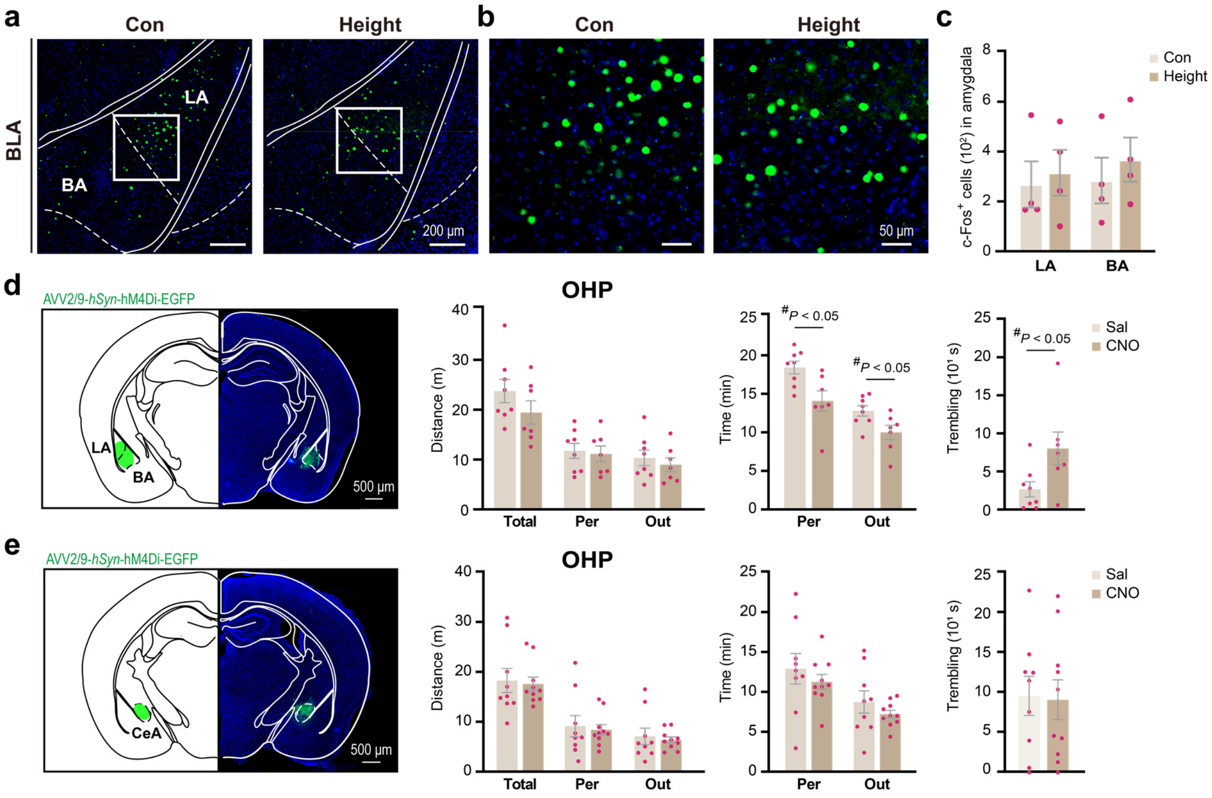

**Fig. S11 Amygdala is dispensable for the fear of heights. a-c** Representative images (**a**, **b**) and quantitative analysis (**c**) of c-Fos signal in different subregions of the amygdala (LA and BA) in response to height exposure. (**d**, **e**) Effects of chemogenetic inhibition of pan-neuronal activities of BLA (**d**) or CeA (**e**) on fear of heights. Data are presented as the mean ± S.E.M. $^{\#}p < 0.05$, a two-tailed Student's *t*-test.

### Supplementary Figure 12

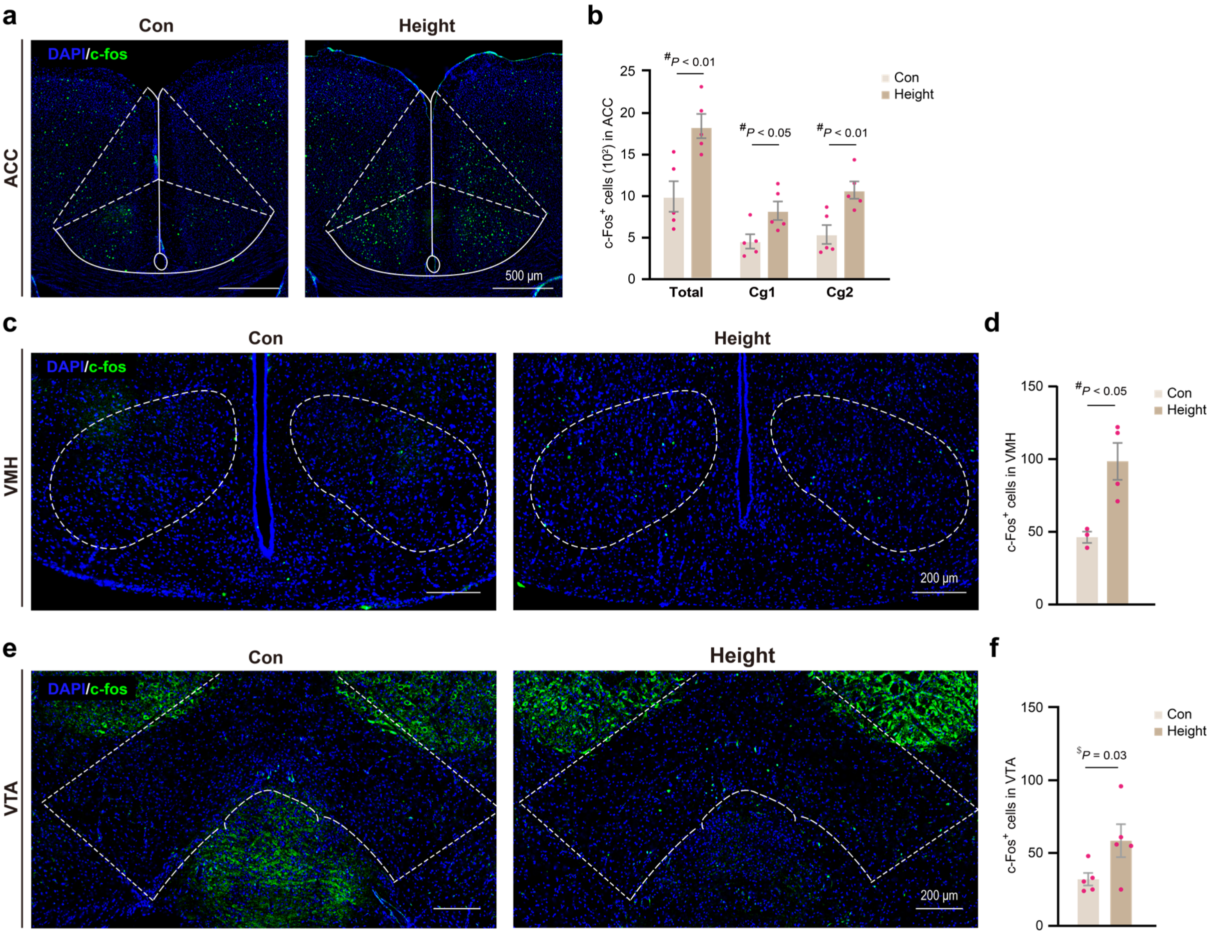

**Fig. S12 Increase in the number of c-Fos+ cells in ACC, VMH, and VTA of mice after height exposure.** Representative images (**a, c, e**) and quantitative analysis (**b, d, f**) of c-Fos+ cells in the anterior cingulate cortex (ACC, **a, b**), ventromedial hypothalamus (VMH, **c, d**), and ventral tegmental area (VTA, **e, f**) of mice in response to height exposure. Data are presented as the mean ± S.E.M. A one-tailed ($) or a two-tailed Student's *t*-test.

**Supplementary Movies**

**Supplementary Movie 1**
Behavior of a mouse on an open high platform (OHP).

**Supplementary Movie 2**
Open high platform (OHP) trembling versus freezing in mice.

**Supplementary Movie 3**
Behavior of a mouse on an elevated platform surrounded by non-transparent walls (GWP).

**Supplementary Movie 4**
Behavior of a mouse on an elevated platform surrounded by transparent walls (TWP).

**Supplementary Movie 5**
Light (left) and dark (right) mouse behavior on an open high platform.

**Supplementary Movie 6**
Swimming of mice with (right) or without (left) gentamicin treatment.

**Supplementary Movie 7**
Wall-climbing behaviors of mice: light, moderate, and heavy.

**Supplementary Movie 8**
Spontaneous nystagmus and tail suspension circling in mice after 24-hour unilateral intratympanic injection of sodium arsanilate.

**Supplementary Movie 9**
Behavioral contrast on OHP between control (left) and l/vlPAG-Inhibited (right) mice.

**Supplementary Movie 10**
Chemogenetic inhibition (middle) and activation (right) of $VGluT2^{+}$ neurons in mouse PAG.
